# PLOD2, a key factor for MRL MSC metabolism and chondroprotective properties

**DOI:** 10.1101/2023.01.18.524662

**Authors:** Sarah Bahraoui, Gautier Tejedor, Anne-Laure Mausset-Bonnefont, François Autelitano, Christian Jorgensen, Mingxing Wei, Farida Djouad

## Abstract

**Background:** Initially discovered for its ability to regenerate ear holes, the MRL mouse has been the subject of multiple research studies aimed at evaluating its ability to regenerate other body tissues and at deciphering the mechanisms underlying it. These enhanced abilities to regenerate, retained in the adult, protect the MRL mouse from degenerative diseases such as osteoarthritis (OA). Here, we hypothesized that MSC derived from the regenerative MRL mouse could be involved in their regenerative potential through the release of pro-regenerative mediators.

**Method:** To address this hypothesis, we compared the secretome of MRL and BL6 MSC and identified several candidate molecules produced at significantly higher levels by MRL MSC than by BL6 MSC. We selected one candidate and performed functional *in vitro* assays to evaluate its role on MRL MSC properties including metabolic profile, migration, and chondroprotective effects. Using an experimental model for osteoarthritis (OA) induced by collagenase (CiOA), we assessed its contribution to MRL MSC protection from OA.

**Results:** Among the candidate molecules highly expressed by MRL MSC, we focused our attention on procollagen-lysine,2-oxoglutarate 5-dioxygenase 2 (PLOD2), coding for the lysyl hydrolase LH2 in charge of post-translational modifications of collagen for its stability and stiffness. PLOD2 is induced by hypoxia-inducible factor 1-alpha (HIF-1a) involved in the regeneration process of adult MRL mice. *Plod2* silencing induced a decrease in the glycolytic function of MRL MSC, resulting in the alteration of their migratory and chondroprotective abilities *in vitro. In vivo*, we showed that *plod2* silencing in MRL MSC significantly impaired their capacity to protect mouse from developing OA.

**Conclusion:** Our results demonstrate that the chondroprotective and therapeutic properties of MRL MSC in the CiOA experimental model are in part mediated by PLOD2.

## Introduction

Regeneration ability is a property that varies widely during development and among species. Indeed, while able to regenerate at early embryonic stages, adult mammals will trigger a tissue repair mechanism resulting in scar formation after tissue injury and limiting the structural and functional recovery [1, 2]. In contrast, some species, such as salamander [3], zebrafish [4] and hydra [5, 6] maintain their regenerative properties during their all-life [7–9]. Rare exceptions, such as the super healer Murphy Roths Large (MRL) mouse, exist among mammals. Indeed, the adult MRL mouse is a competent model for tissue regeneration, suggesting that crucial regenerative mechanisms are conserved in this mammalian model. MRL mouse possesses the extraordinary potential to regenerate multiple musculoskeletal tissues such as the outer ear, articular cartilage, and digits without scarring [10–15]. Among the conserved mechanisms underlying regeneration, the emphasis on aerobic glycolytic energy metabolism has been reported to be essential in several regenerative species [16–18].

MRL mice are protected from the development of joint diseases such as osteoarthritis (OA) [11, 19]. Identifying the mechanisms underlying articular cartilage regeneration and protection from OA would allow the development of novel therapies for OA patients. Indeed, OA is a complex disease characterized, in part, by the degradation of articular cartilage and for which no curative treatment exists to date. One therapeutic option studied is the intra-articular administration of mesenchymal stem/stromal cells (MSC). The trophic activities of MSC introduced exogenously have been shown to protect from cartilage degradation and OA development in experimental model of OA [20–22]. Indeed, such as resident, exogenous MSC participate in joint homeostasis through a paracrine action that contributes to the repair of damaged tissue, the restoration of tissue metabolism and the prevention of inflammation [22, 23].

In the context of digit tip regeneration in mice, mesenchymal cells from one type of tissue were shown to participate to the regeneration of other mesenchymal tissues [24]. The authors showed that the regenerative environment primes the cells from injured tissues to acquire a blastema mesenchymal transcriptional state enabling them to regenerate other mesenchymal tissues such as the dermis. Therefore, it can be speculated that the identification of regenerative environmental factors could stimulate the regenerative potential of MSC in adult mammals. The use of adult MSC in experimental models of degenerative diseases such as OA have shown promising results [20, 25, 26]. However, while MSC prevent cartilage degradation and OA development when administrated locally their capacity to regenerate damaged OA cartilage has never been proven.

The cartilage regenerative potential of MRL mice and their subsequent resistance to experimental osteoarticular defects has increased the interest in MSC derived from MRL mice (MRL MSC). Therefore, we hypothesized that MRL MSC regenerative and protective properties might be associated with their intrinsic properties, particularly their aerobic glycolytic energy metabolism controlled via soluble factors. To address that hypothesis and identify soluble factors presumably at the origin of MRL mouse resistance to OA, we performed a comparative study of MRL MSC and BL6 MSC secretome. Taking into account the glycolytic metabolic profile of the MRL mouse, we investigated the role of one of the genes overexpressed by MRL MSC: *Plod2*, in their regenerative capacities.

## Materials and Methods

### MSC isolation and expansion

Mesenchymal Stem/Stromal Cells (MSC) from MRL/Mpj (MRL MSC) and C57BL/6 (BL6 - MSC) mice bone marrow were isolated and expanded as previously described [27].

### Murine chondrocyte culture and co-culture

Murine articular chondrocytes were isolated from the knees and femoral head of 3-day-old C57BL/6 mice as described previously [28, 29]. Briefly, chondrocytes (25 000 cells/cm^2^) were plated in 12-well culture plates (TPP Techno Plastic Products, Switzerland) with 1 mL of proliferative medium for five days. Then, chondrocytes were treated with 1ng/mL Il-1β (R&D Systems) for 24h (day 0) to generate the so-called “OA-like” chondrocytes. For co-culture experiments, 2 x10^5^ of naïve or modified MSC were seeded in 12-well culture inserts with 1 mL of proliferative media and co-cultured with OA-like chondrocyte for 24h (day 1). 48 hours later, chondrocytes were recovered (day 3) and processed for RT-qPCR.

### Secretome collection and differential quantitative proteome analysis

MSC derived from BL6, BALB/c, DBA1, and MRL mouse bone marrow were cultured in proliferation medium. Then, before being cultured in serum-free DMEM medium without phenol red overnight, the cells were washed in PBS. The conditioned MSC medium was harvested and centrifuged at 3000×g for 5 min at 4°C to remove cells and cell debris and then filtered through a 0.22 μm pore membrane. Supernatants were adjusted to 0.025% with acid labile anionic surfactant I (AALS I, Progenta), concentrated at 4°C in VivaSpin 20 ultrafiltration units (#28-9323-58 MWCO: 3 kDa), and diafiltered at 4°C against 50× volumes of 50 mM NH4HCO3 containing 0.025% AALS I. The protein concentration in the retained material was estimated using the NanoDrop 2000c spectrophotometer (Thermo Fisher Scientific) and scaled to 0.1 mg/mL with diafiltration buffer. Secretomes were stored frozen at −20°C until they were utilized. Label-free quantification and shotgun analysis of secretomes on an Ultimate/Famos/Switchos suite of instruments (Dionex) connected to a hybrid LTQ Orbitrap mass spectrometer (Thermo Fisher Scientific) was performed as previously reported [30, 31].

### MSC transfection with siRNA for PLOD2 silencing

MRL MSC were transfected overnight at subconfluence (45%) with 20mM of control siRNA (siCTL) or the siRNA against PLOD2 (siPLOD2) (Silencer^®^ Pre-designed siRNA, Ambion, Life Technologies™) using Oligofectamine reagent (Life Technologies, Courtaboeuf) according to the supplier’s recommendations. Transfected MRL MSC were used for follow-up experiments 24H post-transfection.

### MSC transfection with CMV plasmid for PLOD2 overexpression

MSC derived from BL6 mice were transfected overnight at subconfluence (60%) with 7,5 ng of PLOD2 plasmid (CMV PLOD2) (pRP[Exp]-mCherry-CMV>mPLOD2 [NM_001142916.1], Vector Builder) using Lipofectamine reagent (Life Technologies, Courtaboeuf) following to the supplier’s recommendations. Transfection level was assessed by using fluorescent microscope (ThermoFisher EVOS™ M5000 Imaging System). Transfected MSC positive for mCherry were used for experiment 24 hours post-transfection.

### RT-qPCR

Total RNA was isolated from mMSC or chondrocytes using the RNeasy Mini Kit (Qiagen, Courtaboeuf), and the quantity and purity of the total RNA were determined using a NanoDrop ND-1000 spectrophotometer (NanoDrop ND, Thermo Fisher Scientific). cDNA was synthesized by reverse transcribing 500ng of RNA into cDNA using the SensiFAST^™^ cDNA Synthesis Kit (Bioline, Meridian Life Science^©^ Company). Quantitative PCR was performed on 6,25ng of cDNA using the SensiFAST^™^ SYBR^®^ No-ROX kit (Bioline, Meridian Life Science^©^ Company) and a LightCycler^®^ 480 Detection system (Roche), following manufacturer’s recommendations. Specific primers for mouse *Plod2*, *Cspg4*, *Inhbb*, *Efemp1*, *Lama4*, *Mmp3*, *Htra1*, *Nt5e*, *Lcn2*, *C1qtnf5*, *Fam20c* and *Hif-1*α were designed using the Primer3 software and can be provided upon reasonable request. Primers for *Col2b, Agn, Mmp13, Adamts5* are the same as previously described [29]. Values were expressed as relative mRNA level of specific gene expression as obtained using the 2^-ΔCt^ method, using the *Rsp9* and *Actin* expression as housekeeping genes.

### Scratch wound healing

Migratory potential of the cells was assessed with scratch wound healing assay. 2.5×10^5^ cells were seeded in TC24 plates and maintained at 37 °C with 5% of CO2 in proliferating media. The wound was performed manually once the cells adhered to the plastic and reached 90% confluency. Wound closure was studied using an inverted microscope (EVOS M5000, Thermo Fisher Scientific) and images of the scratch were taken at H0, just after the scratch and at H24 to evaluate the wound closure. The wounded area was measured at H0 and H24 using Image J Software: the open wound area (in percentage) was calculated by comparing H0 and H24 images and normalized to H0.

### Seahorse assay

Oxygen consumption rate (OCR) and extracellular acidification rate (ECAR) were measured using the XF96 analyzer (Seahorse Biosciences, North Billerica, MA, USA). Transfected and non-transfected murine MSCs (20-25,000 cells/well) were plated on 96-well plates the day before the experiment in XF media (non-buffered DMEM medium, containing 25 mM glucose, 2 mM L-glutamine, and 1 mM sodium pyruvate). The analysis was conducted according to the manufacturer’s recommended protocol. Three independent readings were taken after each sequential injection. The instrumental background was measured in separate control wells using the same conditions without biologic material. The basal glycolytic rate was calculated after glucose injection. The maximal glycolytic rate was measured after oligomycin injection and glycolytic capacity as the difference of oligomycin-induced ECAR and basal stage. OCR was measured under basal conditions and in response to 1 μM of oligomycin, 1 μM of carbonylcyanide-4-(trifluoromethoxy)-phenylhydrazone (FCCP) (Seahorse XF Cell Energy Phenotype Test Kit from Agilent or Sigma Aldrich), and 1 μM of antimycin A and rotenone (Sigma Aldrich). Basal OCR was calculated as the difference between baseline measurements and antimycin A/rotenone-induced OCR; maximal OCR was measured as the difference between FCCP-induced OCR and antimycin A/rotenone-induced OCR. ECAR/OCR ratio was calculated with the glycolytic rate and basal OCR. After the seahorse experiment, the plated cells were fixed for 20 minutes in 4% PFA and then stained with HOESCHT (1/2000eme) to count cells with High-Content Analysis Cellomics™ instrument (Thermo Fisher Scientific) for normalization.

### Collagenase-induced osteoarthritis (CiOA) mouse model and histological analysis

Mice used for this study were housed and cared in accordance with the European directive 2010/63/EU. The CiOA model was generated upon the approval from the Ethical Committee for animal experimentation of the Languedoc-Roussillon and the French Ministry for Higher Education and Research (Approval #5349-2016050918198875). Briefly, 1U type VII collagenase in 5 μL saline was administrated in the intra-articular (IA) space of C57BL/6 mice knee joints (10 weeks old) at day 0 and 2. Groups of 10 mice received MSC (2.5 × 105 cells/5 μL saline) at day 7. At day 42, mice were euthanatized, and paws were recovered for fixation in 4% formaldehyde and decalcified in 4% EDTA solution for three weeks before paraffin embedding. Tibias were sectioned frontally as previously described [20, 29, 30] and stained with safranin O fast green. Two persons performed blind quantification of the degradation of cartilage using the modified Pritzker OARSI score as described [20, 29, 30]. Mice corresponding to uninterpretable stained slides were removed from the analysis.

### Statistical analysis

All data are presented as the mean ± Standard Error of the Mean (SEM), and all experiments were performed at least three times. The Student’s t-test was used to compare two experimental groups, and ANOVA followed by a Friedman test for multi comparison of paired samples was used for the co-culture experiments while ANOVA with Kruskal-Wallis test for multiple comparisons of non-paired samples was used for the CiOA. Graphs show mean ±Standard SEM. P-values < 0.05 (*), P < 0.01 (**) or P < 0.001 (***) were considered statistically significant. Analysis and graphical representation were performed using Graph-Pad Prism^™^ software (Graphpad)

## Results

### MRL MSC exhibit a specific secretome as compared to MSC derived from C57BL/6 mice

We recently performed label-free quantitative shotgun proteomics to identify differentially secreted proteins between MRL MSC and BL6 MSC [30]. We analyzed the secretome of MRL MSC and BL6 MSC based on protein intensities quantified by LC-MS/MS with the aim to identify key factors for MRL MSC metabolism and chondroprotective properties (Fig. 1A). Among the 810 proteins differentially expressed between MRL MSC and BL6 MSC by at least 1.5-fold, 625 proteins were secreted at higher levels by MRL MSC. We focused our attention on proteins with a higher secretory profile by MRL MSC in particular *LAMA4*, *HTRA1*, *PLOD2*, *INHBB*, *MMP3*, *CSPG4*, *NT5E*, *LCN2*, *EFEMP1*, *FAM20C* and *C1QTNF5* (Fig. 1B). By RT-qPCR we confirmed that these 11 factors were overexpressed at a significantly higher level in MRL MSC as compared to BL6 MSC (Fig. 1C).

**Figure 1.**
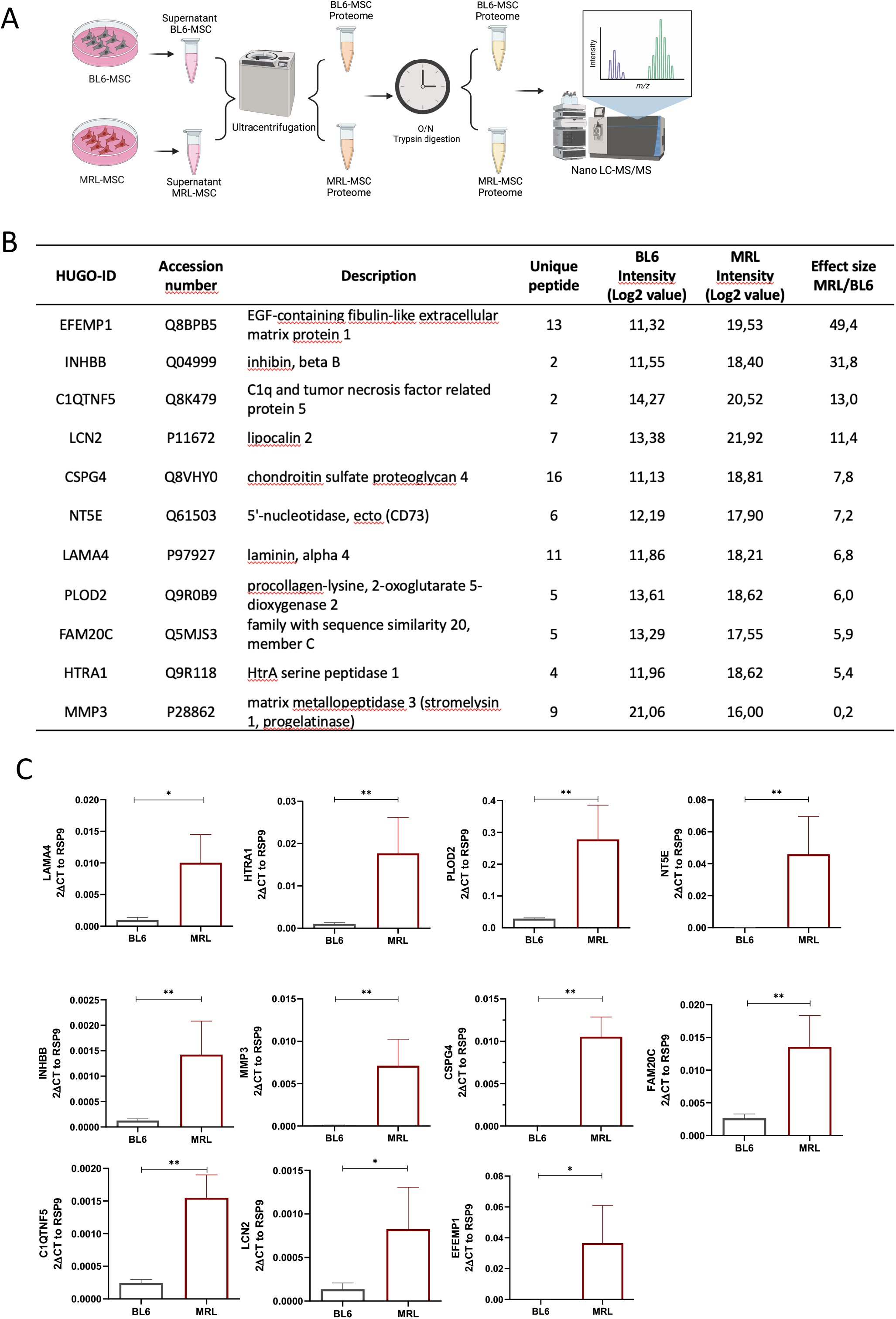
Secretome and expression profiles of MRL MSC and BL6 MSC. (A) and **(B)** Proteomic analysis of differential expression of *Plod2*, *Inhbb*, *Efemp1*, *Lama4*, *Mmp3*, *Htra1*, *Nt5e*, *Lcn2*, *C1qtnf5*, *Fam20c* in MRL MSC compared to BL6 MSC. “Effect size” indicates the standardized mean difference in protein expression level between MRL MSC and BL6 MSC. The median intensity levels (Log 2 value) in MRL MSC and BL6 MSC are indicated for each protein. Normalized protein intensities were used to calculate the Effect size MRL/BL6. **(C)** Increased *Plod2*, *Inhbb*, *Efemp1*, *Lama4*, *Mmp3*, *Htra1*, *Nt5e*, *Lcn2*, *C1qtnf5*, *Fam20c* mRNA levels in MRL MSC compared to BL6-MSC. Error bars represent mean ± SEM. \**P* < 0.05; \*\**P* < 0.01; \*\*\**P* < 0.001, (n=6)

### *Plod2* is required for MRL MSC glycolytic metabolism

MRL mouse use of aerobic glycolysis as their basal metabolic state, characterized by the increased production levels of lactate by cells and in serum, has been proposed [32, 33]. First, we wondered whether MRL MSC exhibit a different metabolism than BL6 MSC. To address that question, we compared the oxygen consumption rate (OCR) and the extracellular acidification rate (ECAR) of the two types of MSC by performing a Seahorse assay (Fig. 2A) We found a significantly lower OCR in MRL MSC (Fig 2B and 2D) while no difference was observed for the ECAR, which can be interpretated as the glycolysis rate (Fig 2C and 2E). The ratio of ECAR to OCR, was significantly higher in MRL MSC than in MSC BL6 suggesting an increased glycolytic function in MSC MRL(Fig. 2F). We then, wondered which factor highly expressed by MRL MSC could be responsible for their glycolytic status. Among the protein overexpressed by MRL MSC as compared to BL6 MSC, we focused our attention on PLOD2 known for promoting aerobic glycolysis in cancer cells [34]. To investigate the role of PLOD2, we used the small interfering RNA (siRNA) approach to knock down the expression of *plod2* in MRL MSC. 48 hours post-transfection of MSC with a siRNA against *plod2* (siPLOD2), *plod2* expression was reduced by 67% compared with the MSC transfected with the control siRNA (siCTL) (Supplementary Figure 1A). We quantified, in MRL MSC either transfected with the siCTL (MRL) or the siPLOD2 (MRL siPLOD2) the OCR, associated to oxidative phosphorylation, and found that *plod2* silencing significantly increased OCR in MRL MSC (Fig. 2G and 2I). ECAR assessment under the same conditions showed that *plod2* silencing did not affect MRL MSCsi PLOD2 when compared to MRL MSC transfection with the control (Fig. 2H and 2J). Overall, the knockdown of *plod2* in MRL MSC induces a switch from glycolysis to OXPHOS as confirmed by the significantly lower ratio of ECAR to OCR in MRL MSC deficient for *plod2* than in MSC transfected with siCTL (Fig. 2K).

**Figure 2.**
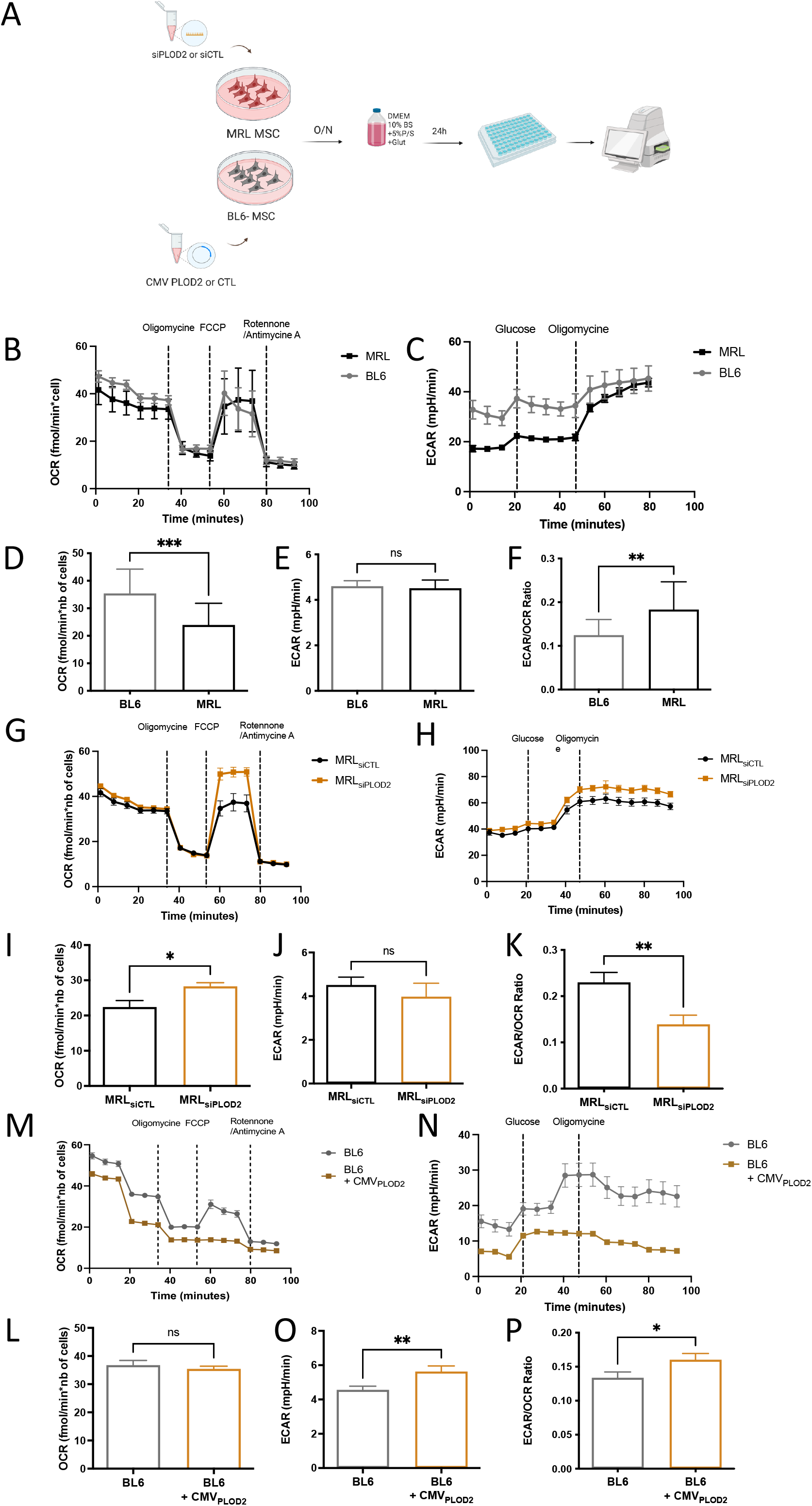
PLOD2 participate in MRL MSC glycolytic metabolism. (A) Workflow of the analysis of OCR and ECAR, performed using Seahorse XF analyzer to assess mitochondrial respiration and glycolysis. **(B)** OCR was measured in BL6 MSC and MRL MSC**, (G)** in MRL MSC_siCTL_ and MRL MSC_siPLOD2_ **and (M)** in BL6 MSC and BL6 MSC_+CM PLOD2_, with sequential addition oligomycin (Oligo, complex V inhibitor), FCCP (a protonophore), and antimycin A (complex III inhibitor)/rotenone (complex I inhibitor) to analyze ATP-linked respiration, basal respiration, maximal respiratory capacity, spare respiratory capacity. (**B, G** and **M**) shows the OCR profiles, and (**D, I,** and **L**) shows mitochondrial bioenergetic parameter calculated from extracellular flux analysis and normalized to the number of cells. (**C)** ECAR was measured in BL6 MSC and (H) MRL MSC in MRL MSC_siCTL_ and MRL MSC_siPLOD2_ **and** (N) in MSC BL6 and MSC BL6_+CM PLOD2_ with serial addition of glucose and oligomycin to measure basal glycolysis, glycolytic reserve, maximal glycolysis. (**C, H,** and **N**) shows the ECAR profiles, and (**E, J,** and **O** shows mitochondrial bioenergetic parameter which can be also interpretated as the glycolytic rate. (**F, K,** and **P**) show the ration ECAR/OCR reflecting the glycolytic capacity. Error bars represent mean ± SEM. \**P* < 0.05; \*\**P* < 0.01; \*\*\**P* < 0.001, Mann-Whitney unpaired t-test, two-tailed (n= 18-32)

Conversely, we asked whether *plod2* overexpression in BL6 MSC would further enhance their glycolytic activity. To address that question, BL6 MSC were transfected with either a plasmid encoding *plod2* (BL6 + CMV_PLOD2_) or an empty vector control (BL6) (Supplementary Figure 1B and 1C) and evaluated the OCR or the ECAR of the cells with Seahorse analyzer. *Plod2* overexpression in BL6 MSC did not modify the OCR (Fig. 2M and 2L) while inducing a switch from OXPHOS to glycolysis as confirmed by the significantly higher ECAR (Fig. 2N-O) and ratio of ECAR to OCR in BL6 MSC overexpressing *plod2* than in control BL6 MSC (Fig. 2P). Altogether, these results provide evidence for the role of *plod2* in the cytosolic glycolytic activity in MSC.

### *Plod2* is required for MRL MSC migration potential

Since a clear relationship between MSC migration and tissue repair has been established [35], we have recently studied the migration potential of MRL MSC and showed that MRL MSC display a significantly higher migration potential than BL6 MSC [30]. To specifically study the role of *plod2* on MRL MSC migratory potential *in vitro*, we analyzed in a scratch wound assay the non-directional migration of MRL and MRL siPLOD2 MSC by evaluating the area of the wound at 24-hours post-wounding using Image J software (National Institutes of Health, Bethesda, MD, USA) (Fig. 3A). Representative images from scratch wound healing assay, 24 hours post-wounding, indicated an altered resurfacing of the wound for MRL siPLOD2 MSC as compared to MRL MSC (Fig. 3B). The percentage of the open wound area at 24 hours which reflects the migration potential of the cells confirmed that MRL siPLOD2 MSC closed the wound significantly slower than MRL MSC (Fig. 3C). Conversely, *plod2* overexpression in BL6 MSC significantly increased the migration potential of BL6 MSC (Fig. 3D) and decreased the open wound area (Fig. 3E). Altogether these results indicate that *plod2* expression plays a positive role on the migration potential MSC.

**Figure 3.**
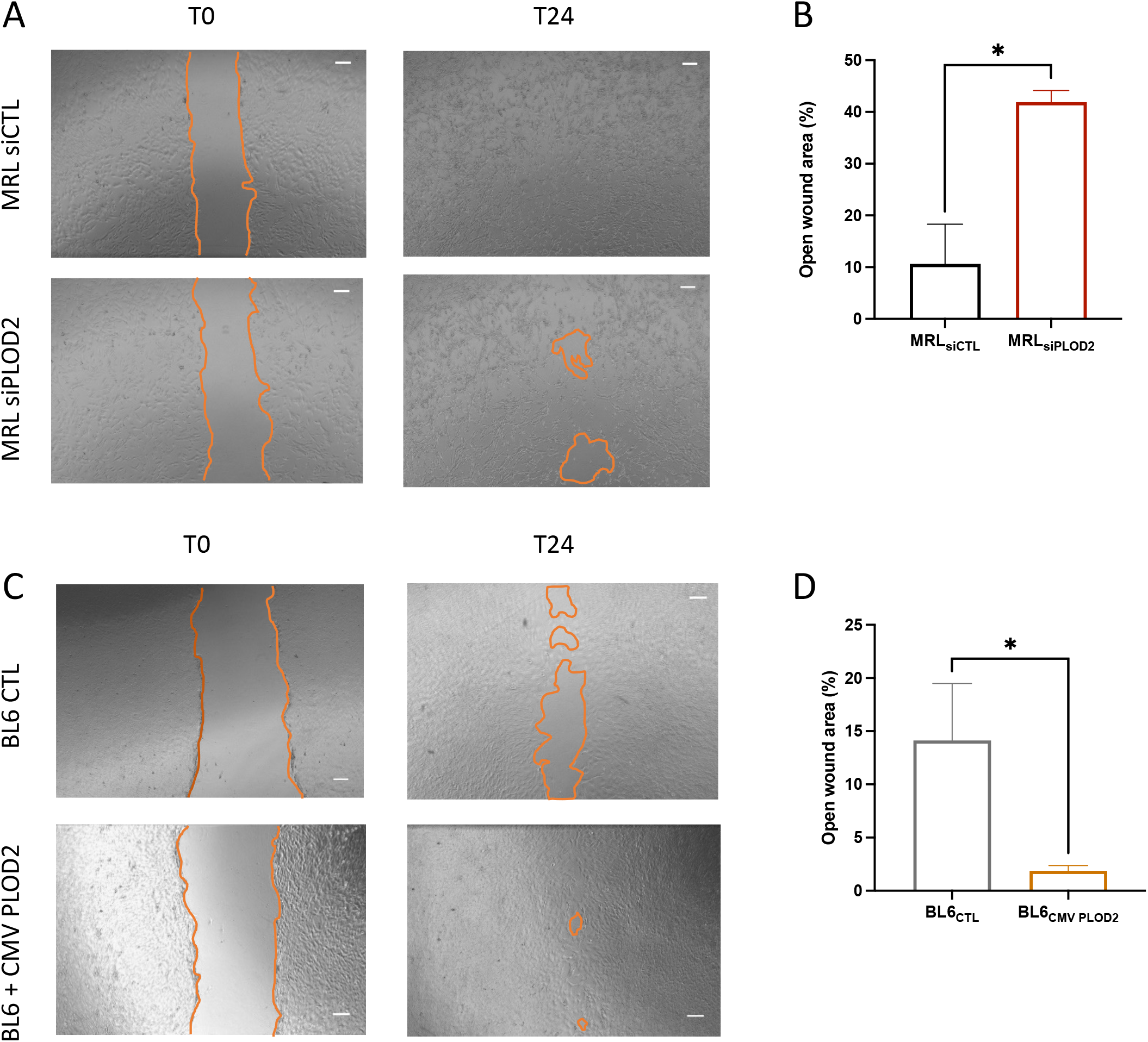
PLOD2 direct MRL MSC migratory ability. **(A)** Workflow (B) Representative images of MSC-MRL_siCTL_ and MSC-MRL_siPLOD2_ and **(D)** BL6 MSC and BL6 MSC_+CM PLOD2_ scratch assay. The images were taken immediately after the scratches had been made and then after 24 h. The orange line indicates the initiatory and final areas without migrating cells. **(C** and **E)** Quantitative analysis of the open wound area was performed at 0 and 24 after wounding using Image J software. 100% corresponds to the highest wound area measured at 0 h. Error bars represent mean ± SEM. \**P* < 0.05; \*\**P* < 0.01; \*\*\**P* < 0.001, Mann-Whitney unpaired t-test, two-tailed (n= 4-5)

### *Plod2* is necessary for MRL MSC chondroprotective properties

MSC protect chondrocytes from the loss of their mature and functional phenotype *in vitro* and *in vivo* [25, 29, 36, 37]. Recently, we have shown that *pycr1*, pivotal for MRL MSC glycolysis, contributed to their pro-anabolic function on chondrocytes [38]. We then wondered whether *plod2* highly produced by MRL MSC could protect chondrocyte from a loss of anabolic markers *in vitro*. To that end, we relied on coculture experiments with MSC and IL-1β-induced chondrocytes in which chondrocytes exhibit a loss of their anabolic markers including *Col2B* and *Acan* and an increase of their catabolic markers and compared the chondroprotective potential of naïve or genetically modified MRL and BL6 MSC (Fig. 4A). First, we tested the effect of *plod2* silencing on MRL MSC chondroprotective effects on the IL-1β-induced chondrocyte model. While co-culture of IL-1β-treated chondrocytes with MRL MSC transfected with a siCtl (mMSC MRL) tend to protect the chondrocytes from a loss of *Col2B* (Fig. 4B), an anabolic marker MRL MSC silenced for *plod2* (mMSC MRL_siPLOD2_) did not (Fig. 4C). Moreover, while co-culture of IL-1β-treated chondrocytes with MRL MSC transfected with a si*Ctl* (MRL MSC) tend to protect the chondrocytes from an increase of ADAMTS5, a catabolic marker, MRL MSC silenced for *plod2* (mMSC MRL_siPLOD2_) did not (Fig. 4C). Conversely, *plod2* overexpression in BL6 MSC (mMSC BL6_CMV-PLOD2_) significantly protect the chondrocytes from an ADAMTS5 increase (Fig. 4D). Altogether, these data show that *plod2* tend to protect MSC from the loss of mature chondrocyte phenotype and the increased expression of catabolic markers which are characteristics of OA.

**Figure 4.**
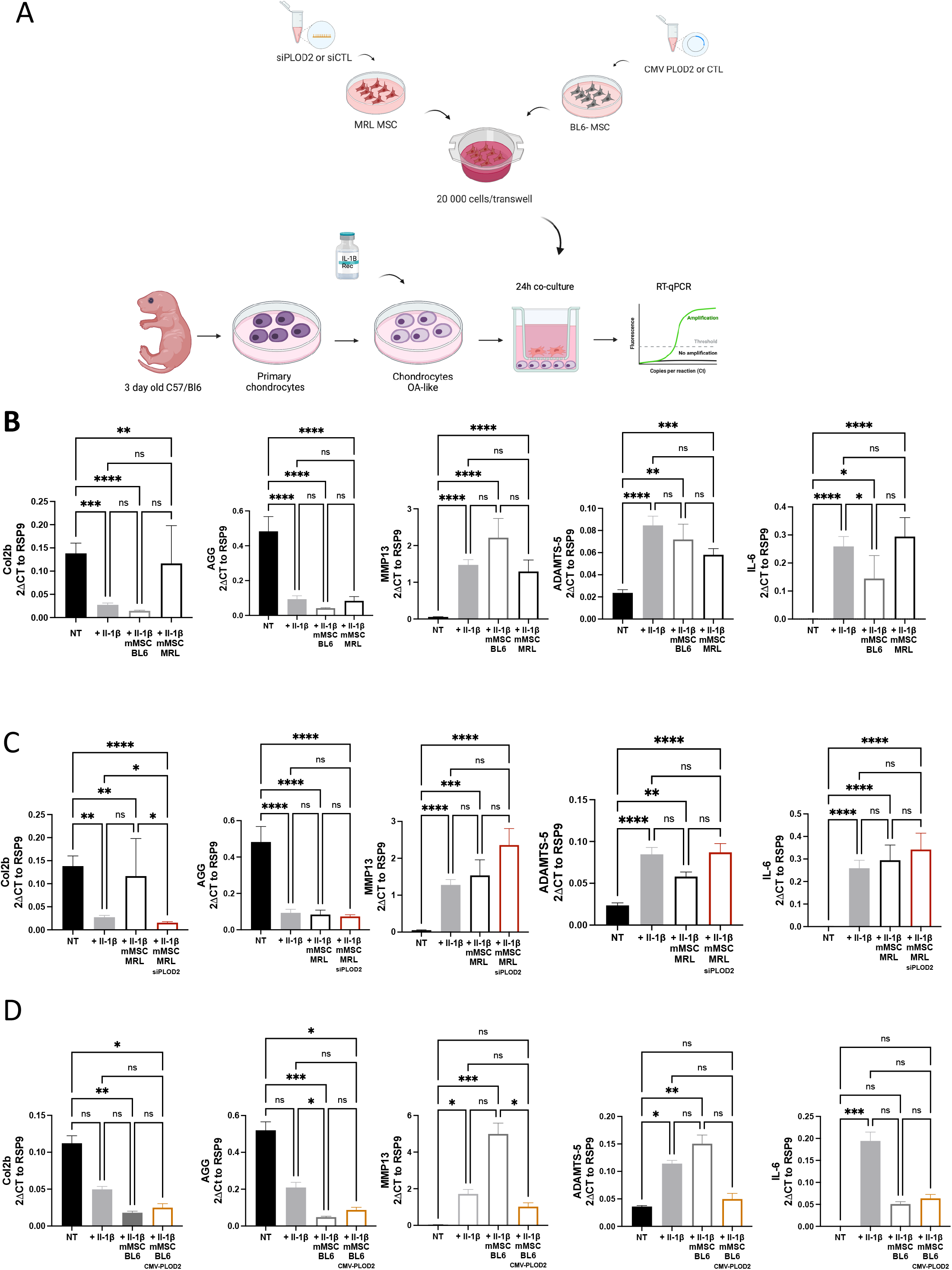
Effect of PLOD2 on chondrocyte gene expression on OA-like chondrocytes. **(A)** Workflow for the generation of OA-like chondrocytes by incubation with IL1β and their co-culture. **(B)** RT-qPCR analysis of different chondrocyte and inflammatory markers in control (NT) and OA-like chondrocytes (IL1β) co-cultured or not with MRL MSC and BL6 MSC (n= 19). **(C)** RT-qPCR analysis of different chondrocyte and inflammatory markers in control (NT) and OA-like chondrocytes (IL1β) co-cultured or not with MRL MSC_siCTL_ and MSC-MRL_siPLOD2_ (n=19). **(D)** RT-qPCR analysis of different chondrocyte and inflammatory markers in control (NT) and OA-like chondrocytes (IL1β) co-cultured or not with MSC BL6 and BL6 MSC_+CM PLOD2_ (n=6). Error bars represent mean ± SEM. One-way paired ANOVA, followed by Friedman test for multiple comparison test was performed. ns: 0.1234; *: p =0.332; **: p=0.0021; ***: p:0;002 or ****: p < 0.0001

### MRL MSC protection against OA is mediated by *plod2*

We subsequently investigated the role of *plod2* on MRL MSC chondroprotective ability *in vivo*. For this purpose, we used the collagenase-induced osteoarthritis (CiOA) model, in which mice show signs of cartilage degradation, to test the effect of intra-articular injection of MRL MSC transfected with siCTL (MSC MRL) or siPLOD2 (MSC MRL_siPLOD2_). At D42, histological examination revealed a lower osteoarthritis score in collagenase-treated mice injected with MRL MSC than in collagenase-treated mice without MRL MSC, indicating a protective effect of the MRL MSC (Figure 5A and 5B). In contrast, the OA score of mice injected with MRL MSCsilenced for *plod2* (MSC MRL_siPLOD2_) was significantly higher than that of untreated mice (injected with PBS) or CiOA mice treated with control MRL MSC (Figure 5A and 5B). Those results suggest that *plod2* contributes to the protective effect of MSC MRL against OA.

**Figure 5.**
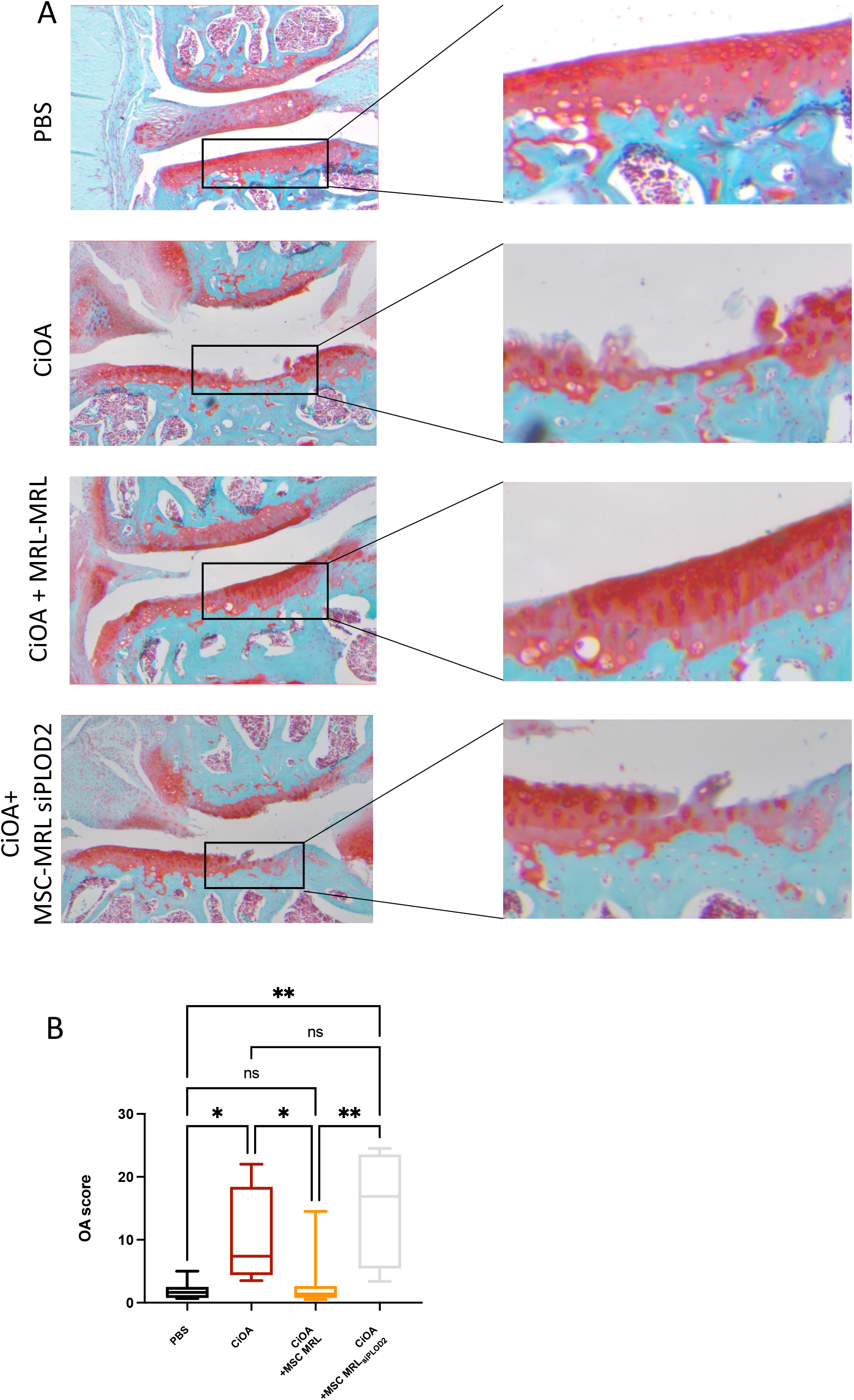
*Plod2* mediates MRL MSC chonrdroprective effect from osteoarthristis. **(A)** Histological sections of CiOA mice not treated (PBS), treated with collagenase only (CiOA), collagenase, MRL MSC (CiOA + MSC MRL) and MRL MSC With siRNA for PLOD2 (CiOA+MSC MRL_siPLOD2_) **(B)**. OA score of histological sections of knee joints of the mice described in **(A)** Error bars represent mean ± SEM. One-way ANOVA, (Kruskal-Wallis test) was performed. ns: 0.1234; *: p =0.332; **: p=0.0021; ***: p:0;002 or ****: p < 0.0001. n=8 for PBS, CiOA an CiOA+ MSC MRL_siPLOD2_and n=9 for CiOA + MSC MRL

## Discussion

Unlike most mammals, the “super healer” MRL mouse retains its regenerative capacity in adulthood. Indeed, this mouse can regenerate, among other tissues, its muscles, its nervous system, and its cartilage. This particularity could explain why this animal model does not develop specific degenerative pathologies such as osteoarthritis or osteoporosis when induced experimentally. To better understand their resistance to joint degenerative diseases we focused our attention on the secretome of their MSC in comparison with BL6 MSC and demonstrate a specific secretome of MRL MSC with a significantly higher production of PLOD2 compared to BL6 MSC. *Plod2* produced by MRL MSC assigns them their glycolytic status as well as their chondroprotective properties.

PLOD2, involved in lysyl hydroxylation of collagen molecule pivotal for the stability of collagen cross-links [39], is regulated by hypoxia-inducible factor (HIF)-1α [40, 41]. While the majority of research investigating the contribution of PLOD2 in tumorigenesis [42, 43], our study provides the first evidence of the role of PLOD2 on MRL MSC metabolism, migration potential and chondroprotective properties. HIF1α plays a crucial role in MSC functions [44] and several if not all mammalian regeneration processes [45]. In MRL mice, after tissue wounding, the biphasic expression profile of HIF1α, characterized by a rapid increase of systemic HIF1α levels peaking between days 10 and 14 and decreasing during the second of the regeneration process, suggested the critical role of HIF1α during the process [45, 46]. This was confirmed in experiments showing that *hif-1α* silencing in MRL mice inhibited ear hole closure and that the injection of drugs stabilizing HIF1α in a hydrogel both proximal and distal to the injured sites led to an accelerated ear hole closure [45, 47]. Since HIF1α is a regulator of PLOD2 expression [48] [49], we assessed the expression level of *hif-1α* in MRL MSC silenced for *plod2* and found that *hif-1a* was reduced in MRL MSC transfected with the siRNA against *plod2* as compared to MRL MSC transfected with the siCTL (Supplementary Fig. 2A and 2B). Conversely, *plod2* overexpression in BL6 MSC causes a metabolic switch from oxidative phosphorylation to glycolysis. Altogether, these results suggest that in MRL MSC PLOD2 regulates *hif-1α* expression and that the PLOD2-HIF1α axis controls MRL MSC glycolysis and regenerative properties we have described [38] and recent findings in the literature showing the induction of PLOD2 by the HIF1α axis for ECM remodeling [41].

*In vitro*, in gain and loss of function experiments, we have shown that *plod2* expression increases MRL MSC migration. These results are in line with a study showing that the inhibition of hypoxia-induced PLOD2 reduces the migration and the invasion of glioma cell both *in vitro* and *in vivo* [50]. This was associated with an elevated expression of E-cadherin and reduced expression of *vimentin, N-cadherin, snail* and *slug* in response to PLOD2 suppression. Further experiments should be performed to determine whether the decreased migration potential of MRL MSC in response to *plod2* silencing is due a modulation of adhesion molecule expression levels.

The silencing of *plod2* in MRL MSC also altered MSC chondroprotective properties on IL-1β-treated chondrocytes. Indeed, we demonstrated that while MRL MSC protects IL-1β-treated chondrocytes from a loss of anabolic marker *Col2B*, MRL MSC silenced *plod2* did not. These results are consistent with our results in the CiOA model where MSC MRL downregulated for *plod2* lose their chondroprotective effect. Therefore, we propose that the chondroprotective effect of MRL MSC relies, in part, on *plod2* overexpression.

Our results are in contradiction with Bank et al. study suggesting that PLOD2 inhibition and therefore the prevention of the formation of pyridinoline cross-links which stabilize the collagen and, might favor cartilage repair attempts. They argue that by showing cartilage with collagen-containing low levels of hydroxylysine and pyridinoline might be less prone to degradation induced mechanically [51]. Therefore, the positive role of *plod2* expression on MRL MSC chondroprotective effect might be due to other properties of *plod2* than that to form collagen cross-links. Stegen and colleagues recently showed that collagen synthesis in chondrocytes was metabolically controlled by *hif-1α* and *plod2* post-translational modifications [52]. However, sustained expression of *plod2* can lead to bone dysplasia, suggesting its involvement in fibrosis [52]. Interestingly, overexpression of PLOD2 via the TGF-1 pathway in adipose tissue-derived MSC increases the therapeutic potential of MSC in an experimental model of spinal cord injury [53]. In mice with a dominant-negative mutation of the TGF-type II receptor, disorganization of collagen fibers was observed [54]. In view of the discrepancy in these results, it would be interesting to know whether the TGF pathway is also used by the MRL mouse for PLOD2 induction and whether this is beneficial against OA.

We have recently shown that the chondroprotective effect of MRL MSC has been associated with their glycolysis [38]. Indeed, we showed that Pyrroline-5-Carboxylate Reductase 1 (*Pycr1*) downregulation induced MRL MSC metabolism reprogramming specifically associated with a reduced lactate concentration in the extracellular media of the cells. This OXPHOS metabolic reprogramming of MRL MSC knockdown for *Pycr1* induced a loss of MRL MSC chondroprotective functions. It is thus reasonable to propose that PLOD2 is pivotal for MRL MSC chondroprotective functions through the high glycolytic flux produced by MRL MSC.

## Conclusions

In conclusion, our findings demonstrate for the first time that the enhanced chondroprotective potential of MRL MSC is attributed, in part, to PLOD2, which controls their higher glycolytic potential compared to BL6 MSC.

## List of abbreviation

CiOA: Collagenase-induced osteoarthritis
MRL: Murphy Roths Large
MSC: Mesenchymal stromal/stem cells
ECAR: extracellular acidification rate
HIF-1a: hypoxia-inducible factor 1-alpha
OA: osteoarthritis
OCR: Oxygen consumption rate
PLOD2: procollagen-lysine,2-oxoglutarate 5-dioxygenase 2
RT-PCR: reverse transcription-polymerase chain reaction
SEM: Standard Error of the Mean

## DECLARATIONS

### Ethics approval

The CiOA model was generated upon the approval from the Ethical Committee for animal experimentation of the Languedoc-Roussillon and the French Ministry for Higher Education and Research (Approval #5349-2016050918198875).

### Consent for publication

« Not applicable »

### Availability of data and material

The datasets during and/or analyzed during the current study available from the corresponding author on reasonable request.

## Acknowledgements

We thank the MRI facility for their assistance and Daouda Abba Moussa from Metamontp for his help in the Seahorse experiment. We thank the Réseau d’Histologie Expérimentale de Montpellier (Plateau RHEM) histology facility for tissue processing and the animal facility RAM-Neuro for his help in animal breeding and handling. All the illustrations were created with Biorender.com

## Author Contributions

FD and SB designed the whole project and the experiments with input of FA, GT. SB performed the experiments and analysed the results with the input of GT, A-L.M-B and FA. FD and SB wrote the manuscript with the input of CJ, MW, FA. All authors read and approved the final manuscript.

## Figure Legends

**Supplementary figure 1. siRNA and plasmid transfection control**

**(A)** RT-qPCR analysis of *Plod2* in MRL MSC transfected with control (siCTL) or anti-PLOD2 siRNA (siPLOD2) 24h and 48h post-transfection (n=1) (**B)** RT-qPCR analysis of *Plod2* in BL6 MSC transfected with plasmid CMV PLOD2-mCherry (BL6+CMV PLOD2). **(C)** BL6 MSC transfection with CMV PLOD2 was assessed for mCherry expression.

**Supplementary figure 2. *Hif-1a* mRNA levels in MRL MSC**

**(A)** RT-qPCR analysis of *Hif-1a* in MRL MSC transfected with control (siCTL) or anti-PLOD2 siRNAs (siPLOD2) 24h post-transfection (n=1) **(B)** RT-qPCR analysis of *Hif-1a* in BL6 MSC and MRL MSC Error bars represent mean ± SEM. \**P* < 0.05; \*\**P* < 0.01; \*\*\**P* < 0.001, Mann-Whitney unpaired t-test, two-tailed (n=3

