## Supplementary figures and images for "PLOD2, a key factor for MRL MSC metabolism and chondroprotective properties"

### Supplementary Figures 1 and 2

##### Supplementary Data 1

A

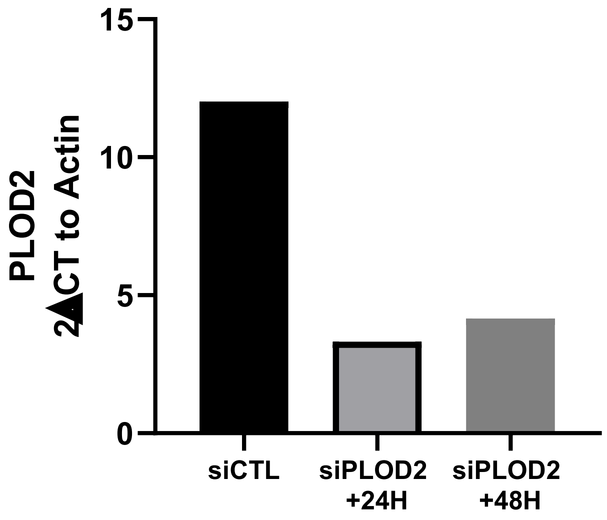


B

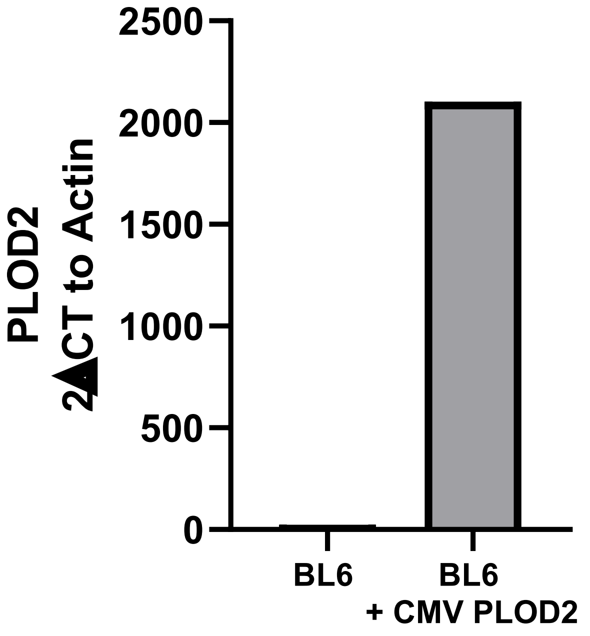


C

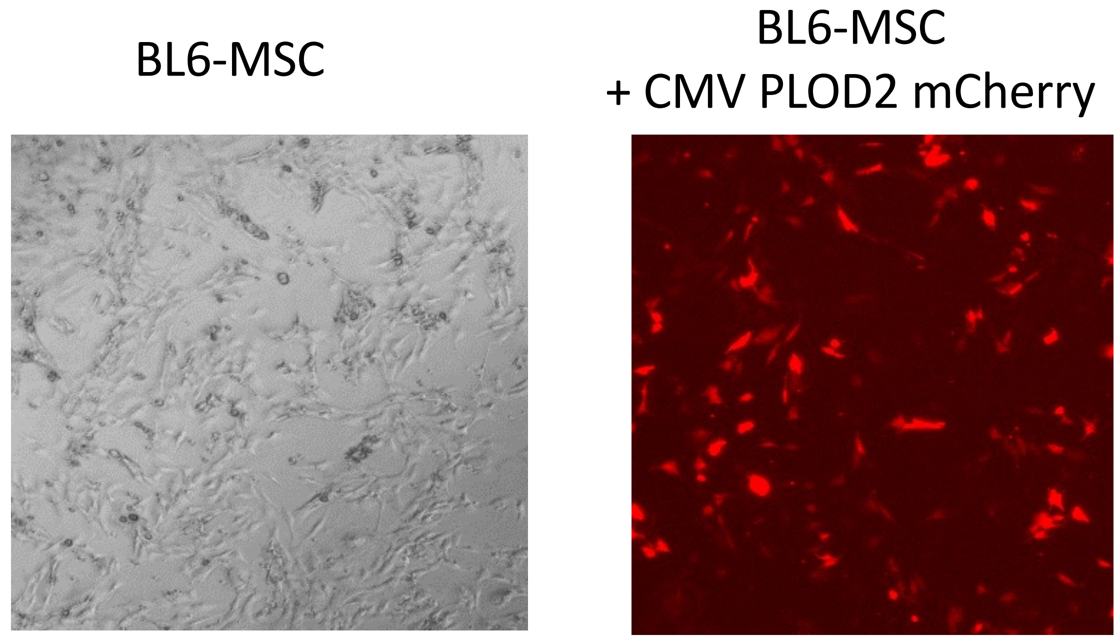

#### Supplementary data 2

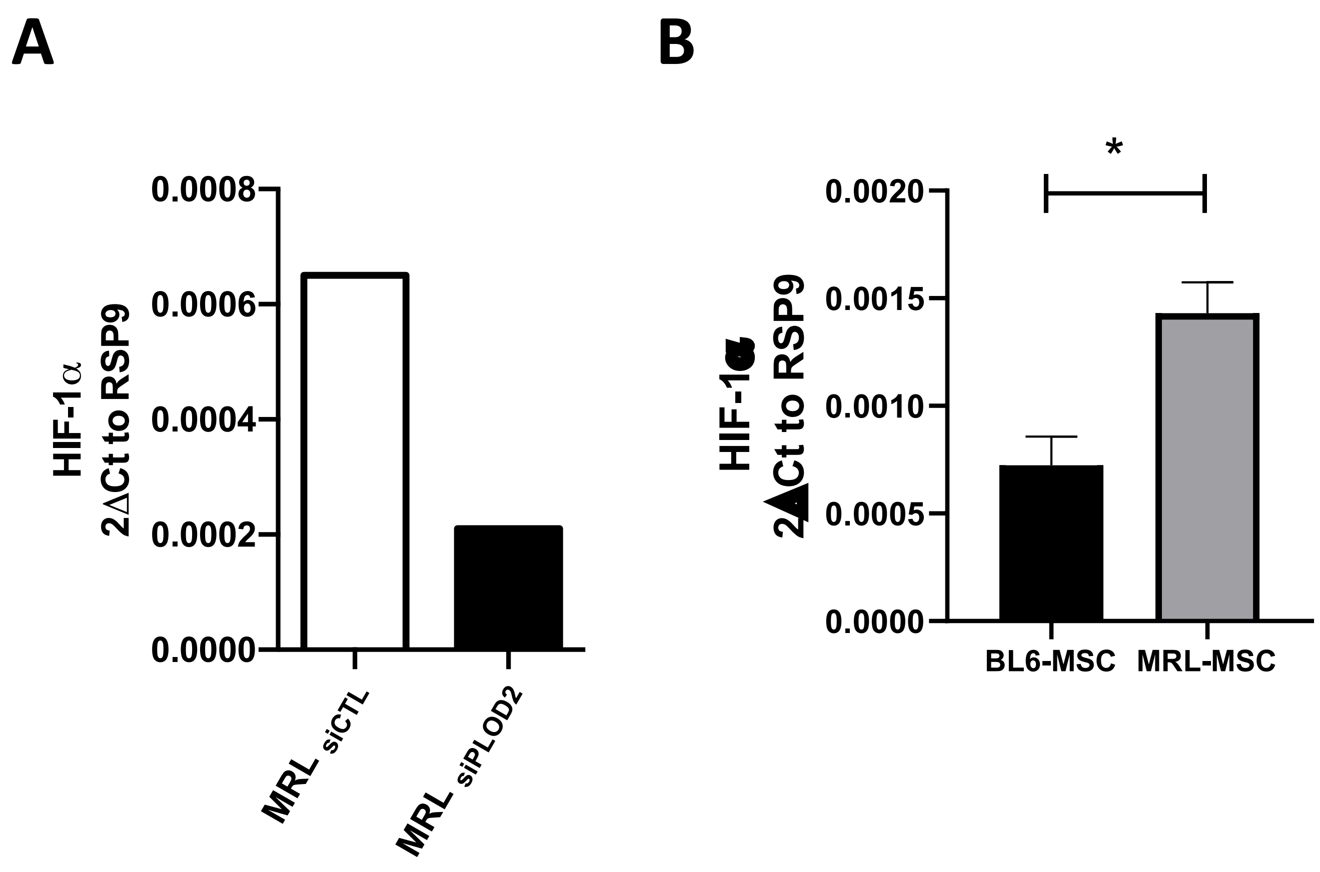
